# Neural Signatures of Predictive Processing during Natural Mandarin Speech Comprehension

**DOI:** 10.1101/2025.11.23.690006

**Authors:** Qifei Wang, Jakub Szewczyk, Judit Fazekas, Eva Berlot, Floris P de Lange

## Abstract

Language comprehension requires the continuous transformation of speech into a hierarchy of linguistic units, from phonemes to syllables to words. Because speech unfolds rapidly, listeners are thought to predict upcoming content to keep pace. Previous research has provided empirical evidence for predictive processes operating at multiple linguistic levels during naturalistic listening, including words and phonemes. However, it remains unclear whether prediction also operates concurrently at other levels, such as syllabic and phrasal representation. Here we use Mandarin Chinese to examine the neural signatures of predictive processing across multiple levels of linguistic granularity during natural speech comprehension. Mandarin comprises four representational levels: phoneme, sub-syllabic, character and word, and its lexical identity is largely constrained at the sub-syllabic level, potentially redistributing predictive weight across linguistic representations. We recorded magnetoencephalography (MEG) data while 34 native Mandarin speakers (21 females) listened to a naturalistic audiobook and applied linear regression modeling to examine how linguistic features modulated neural activity. We found that the brain activity of listeners segmented speech into hierarchical units, and that surprisal modulated responses simultaneously across sub-syllabic, character and word levels. In contrast to findings from Indo-European languages, however, we did not observe unique surprisal effects at the lowest, phonemic level. Furthermore, the surprisal of lexical tone in Mandarin modulated brain activity only when integrated with its phonological components. These findings suggest that predictive processing during Mandarin speech comprehension operates concurrently across multiple (though not necessarily all) levels of linguistic granularity, with its implementation shaped by language-specific structural properties.

**Significance statement:** Language comprehension involves segmenting a continuous acoustic stream into multiple linguistic units, from phonemes to words, and generating predictions at these levels. However, direct neural evidence remains limited regarding how segmentation and prediction operate simultaneously across levels of linguistic granularity, particularly outside Indo-European languages. Using temporal response function analysis of magnetoencephalography data recorded during naturalistic Mandarin listening, we show that predictive processing occurs across multiple levels of linguistic granularity. Specifically, we find evidence for prediction-related neural responses at sub-syllabic, character, and word levels, but not a reliable unique effect at the phonemic level. These results indicate that predictive processing also operates during Mandarin speech comprehension, and its neural implementation is shaped by language-specific structural properties.

## Introduction

When listening to speech, the brain performs a remarkable feat: it transforms a continuous acoustic stream into a structured and meaningful message. This process begins with the segmentation of the incoming signal into discrete linguistic units–such as phonemes, syllables, words, and phrases (Brent, 1999; Mattys et al., 2005; Sanders & Neville, 2000). The brain tracks the time courses of these abstract structures in parallel (Ding et al., 2016; Giraud & Poeppel, 2012) and interprets them incrementally as speech unfolds (Marslen-Wilson, 1973). As natural speech is rapid (Crystal & House, 1990) and often degraded by noise (McDermott, 2009), it is thought to be essential for the brain to actively predict upcoming content. Such predictive processing enables the listener to disambiguate the signal in real time and maintain pace with the speaker without a delay (Clark, 2013; Pickering & Garrod, 2013).

Substantial evidence supports predictive mechanisms and their implementation across multiple linguistic levels (Heilbron et al., 2022; Kuperberg & and Jaeger, 2016; Ryskin & Nieuwland, 2023). For instance, while listening to a speech fragment (e.g., the boy went out to fly a …), the brain actively predicts not only the upcoming word (’kite’; Weissbart et al., 2020; Willems et al., 2016), but also the next phoneme (’/k/’; Brodbeck et al., 2022; DeLong et al., 2005; Donhauser & Baillet, 2020) based on prior context. However, although speech is represented across multiple levels of linguistic granularity (Christiansen & Chater, 2016; Keitel et al., 2018), direct neural evidence remains limited regarding how predictive processing is distributed across these levels during natural speech comprehension. At the same time, although phoneme-level prediction has been reported in prior studies, it remains unclear whether phonemes play an equally central role in languages that organize linguistic structures around additional or alternative units.

Mandarin is particularly suitable for investigating these questions because it naturally defines four levels of linguistic representation (**Fig. 1**): *phonemes*, the smallest contrastive units that can alter lexical meaning (Chomsky & Halle, 1968); *sub-syllabic units*, which can be composed of multiple phonemes, and serve as a crucial instructional tool in Chinese literacy education; *characters*, the basic units of the writing system that typically correspond to both a syllable and a morpheme; and *words*, which are composed of one or more characters that convey complete meaning. This hierarchical structure offers a clear framework to examine how segmentation and prediction unfold across different linguistic levels. Moreover, phonemes in Mandarin play a relatively reduced role in determining lexical identity compared to many Indo-European languages (Yi & Duanmu, 2015). Because the inventory of phonemic combinations within sub-syllabic units is limited, individual phonemes often carry less distinctive information in signaling word meaning. At the same time, predictive processing is thought to operate over stored representational units and to be shaped by the statistical structure of a language (Huettig et al., 2022; Kuperberg & Jaeger, 2016). Consequently, predictive processing in Mandarin may rely more strongly on sub-syllabic representations than on individual phonemes.

Another fundamental and distinctive phonemic feature of Mandarin is lexical tone, which distinguishes meaning among syllables with identical segmental structures. For example, /ta1/ means 她 (“she”), whereas /ta3/ means 塔 (“tower”). Evidence for tonal prediction in Mandarin remains mixed, with some object-naming paradigms showing no tone-specific prediction (Xu et al., 2024; Z. Zhao et al., 2024), while other studies have revealed decodable neural signal reflecting tone prediction (Liu et al., 2025). Nevertheless, these findings derive from controlled and artificial paradigms, leaving open the question of whether tone-related predictability modulates neural responses during naturalistic speech comprehension.

In the present study, we analyzed magnetoencephalography (MEG) recordings from native Mandarin speakers listening to a Chinese audiobook. We applied time-resolved multivariate temporal response function (mTRF) analysis (Crosse et al., 2016) to characterize the temporal dynamics of neural responses to multiple linguistic features as the narrative unfolded. Onset-related neural responses were used to examine how continuous speech is segmented into discrete linguistic units, as onset-evoked responses are reliable markers of speech segmentation (Sanders et al., 2002). Predictive processing was assessed using surprisal, quantified with a Chinese GPT-2 model in a fine-grained, context-sensitive manner. Surprisal is commonly used as a proxy for prediction error, with higher surprisal reflecting a greater mismatch between context-based expectations and actual input (Kuperberg & Jaeger, 2016; Levy, 2008; Shain et al., 2024). Words with high surprisal typically impose greater processing demands and elicit stronger neural responses, particularly around 400 ms after word onset (Kutas & Federmeier, 2000; Kutas & Hillyard, 1980).

**Figure 1.**
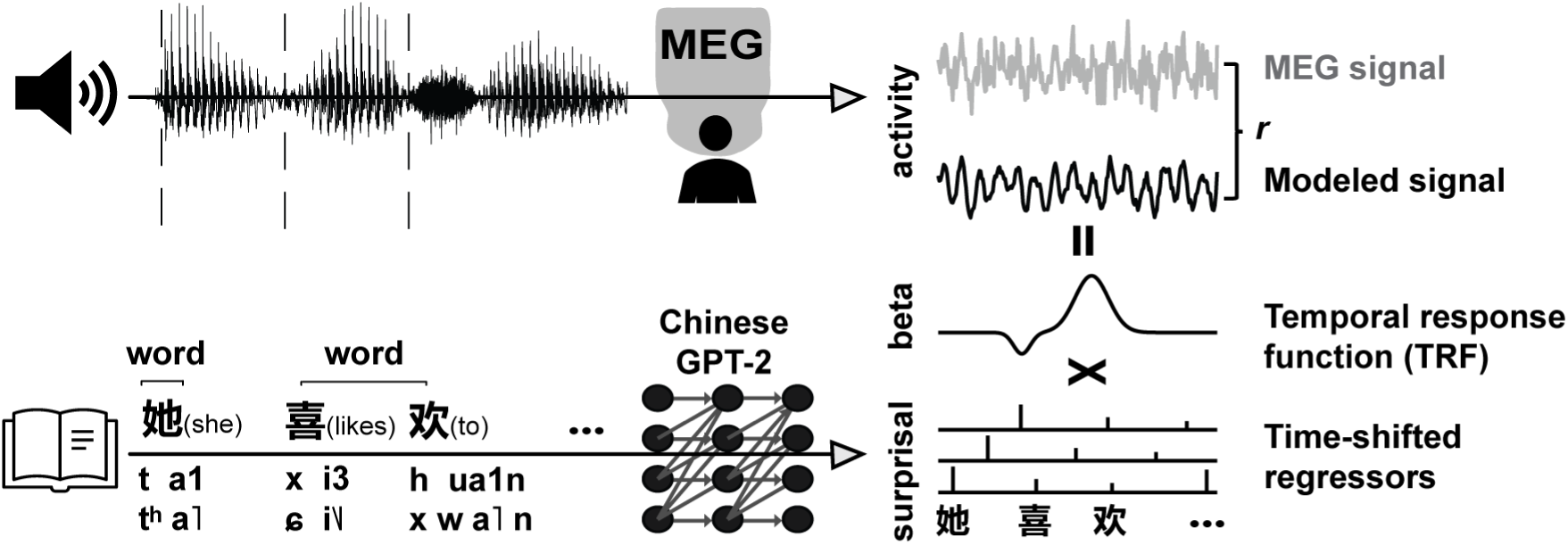
Experimental paradigm and analysis pipeline. Participants listened to a Chinese audiobook while their brain activity was recorded using magnetoencephalography (MEG). The linguistic structure of Mandarin is illustrated: words comprise one or more characters; each character maps to a single syllable, which is composed of sub-syllabic units—an optional *initial* (consonant) and a *final* (vowel(s), sometimes with a nasal consonant). The final also carries a lexical tone (indicated by numbers 1-4). Sub-syllabic units are further composed of one or more phonemes. The audiobook transcript was analyzed using a Chinese GPT-2 model to quantify contextual surprisal for each character. Temporal response function (TRF) modelling was used to estimate the linear relationship between linguistic features and MEG signal. As illustrated for character-level surprisal, the regressor was time-shifted (steps of 1/120 s, spanning from 100 ms before to 1000 ms after character onset) to construct time-lagged regressors. TRF model estimates the beta weights at each time lag, capturing the unique contribution of this surprisal regressor to the MEG signal. Model performance was evaluated via cross-validation, calculating the Pearson’s correlation (*r*) between the predicted (black) and observed (grey) neural responses on held-out data.

## Materials and methods

### Participants

Thirty-four native speakers of Mandarin Chinese were recruited (female = 21, aged from 21-36 years, mean = 28 years, SD = 3.73 years). All participants were native speakers of standard Mandarin and were fluent in English. None reported any neurological, developmental, or language impairments. All participants were right-handed, with normal hearing and normal or corrected-to-normal vision. They provided written informed consent for their data to be used for research purposes and received monetary compensation for their participation. The study was approved by the local ethics committee (CMO 2014/288; CMO Arnhem-Nijmegen, the Netherlands) and conducted in accordance with the Declaration of Helsinki. The sample size was determined based on a priori power analysis (Faul et al., 2009), which indicated a sample of 34 would achieve 0.8 power to detect a medium effect size (Cohen’s *d* = 0.5) at ɑ = 0.05 (two-tailed paired *t* test).

### Stimuli and procedure

All audio stimuli were presented using Matlab Psychtoolbox (Brainard, 1997; Kleiner et al., 2007) and played through a loudspeaker positioned directly in front of the participant at a clear and comfortable listening volume. Participants sat comfortably while their brain activity was continuously recorded using magnetoencephalography (MEG).

The listening material consisted of the opening chapters of the audiobook *Bailuyuan*, split into 40 segments ranging from 56 to 91 seconds in duration (totaling 50.7 minutes and 9,945 characters). All recordings were presented during a single experimental session. Segments were presented in chronological order to preserve contextual continuity.

Following each segment, participants performed a brief recognition task. They heard two short audio clips (each ∼3 seconds), one from the segment they had just heard and the other from a later, unused portion of the audiobook. The two clips were presented in a random order, and participants indicated which one had appeared in the preceding segment using a button press. Four buttons were used and responded to with the index and middle fingers of each hand: the left hand indicating the first clip and the right hand the second. Confidence was reported by the exact finger: middle fingers indicated high confidence and index fingers low confidence. Feedback on the correctness of participants’ choice was provided after each response. The task was intentionally simple and served primarily to ensure sustained attention during listening (mean accuracy = 95.3%). Note that these periods of recognition task with response and feedback were excluded from the MEG analysis. After each trial, participants could take a break and start the next segment at their own pace.

### Stimuli annotation

To obtain precise onset times for each character and sub-syllabic unit, the manually transcribed text was aligned with the audio using the Montreal Forced Aligner (MFA; McAuliffe et al., 2017). This alignment provided onset and offset timestamps for each character and each phoneme. All automatically generated onsets and offsets were then visually inspected and manually adjusted in *Praat* (Boersma & Weenink, 2011) to ensure temporal accuracy.

Word boundaries were determined using HanLP (He & Choi, 2021) toolkit for word segmentation, which was then visually inspected for correction, resulting in 5880 words. Word onsets were defined as the start time of the word’s first character. Each character was converted to its phonological form using the *pinyin* Python package (Huang et al., 2025). The sub-syllabic structure of each character was then parsed: the initial (where present) was treated as the first sub-syllabic unit, and the final with tone as the second. The onset times for all sub-syllabic units were assigned based on phoneme-level alignment obtained from MFA.

### MEG acquisition and preprocessing

Brain responses were recorded with a whole-head 275-channel axial gradiometer MEG system (CTF) inside a magnetically shielded room at the Donders Centre for Cognitive Neuroimaging in Nijmegen, the Netherlands. MEG signals were digitized at a sampling rate of 1200 Hz with an online 300-Hz low-pass filter applied during acquisition. Participants’ head positions were continuously monitored using localization coils placed at the nasion and the left and right pre-auricular points. If head movement exceeded 5 mm from the initial position, participants were asked to readjust it during breaks. In addition, participants’ left eyes were tracked using an SR Research Eyelink 1000 eye tracker, and behavioral responses were collected using MEG-compatible button boxes.

MEG data were preprocessed and analyzed using the Fieldtrip toolbox in Matlab (version 2023b; Oostenveld et al., 2011). Continuous recordings were segmented into 40 trials based on the timestamps of the 40 audio segments. A notch filter (Butterworth IIR) was applied at 49-51, 99-101, and 149-151 Hz to remove potential line noise artifacts. Next, data was band-pass filtered between 0.1 - 20 Hz using a bidirectional finite impulse response filter to focus on slow, evoked responses (Heilbron et al., 2022).

Data were then downsampled to 120 Hz and cleaned for artifacts such as muscle contractions and SQUID jumps using a semi-automated approach. Specifically, trials were divided into 500 ms snippets with 200 ms overlap between them. Snippets containing abnormally high variance were identified through visual inspection and removed using Fieldtrip function *ft_rejectvisual* with the ‘summary’ method. On average, 0.39% of the data were removed per participant (ranging from 0.01% to 2%). The remaining snippets were then combined to reconstruct the original 40 trials and demeaned within each trial to remove any DC drift. Independent component analysis (ICA) was subsequently applied to remove residual artifacts associated with eye movements, cardiac activity, and other non-neural noise sources.

### Regressors of linguistic features

#### Onset regressors

Onset regressors were constructed as impulse functions with a value of 1 at the onset of each language unit–phoneme, sub-syllabic unit, character and word, with values being 0 elsewhere. Each regressor spanned the entire length of the neural signal.

#### Regressors of prediction

Linguistic prediction was estimated using Chinese GPT-2 (Fig 1; Zhao et al., 2019), a large pretrained language model that performs comparably to humans in next-word prediction tasks. The model’s basic token is the character. The text of the audiobook was passed through GPT-2 to obtain conditional probability of each character based on its preceding context.

To determine the optimal context window for probability estimation, we generated predictions using windows of 1, 2, 3, 8, 16, 32, 64, 128, 256, 512 and 1024 tokens. The 1024-character window provided the best model fit and was therefore used for all subsequent analyses (**Fig. 3D**). These character-level probabilities were used to derive probabilities for words, sub-syllabic units, phonemes, tones, and rhymes. For a word W composed of characters {c_1_,…,c_n_}, the word probability was defined as the joint probability 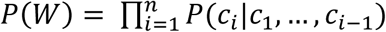 For phonological features (e.g., phonemes, sub-syllabic units, tones, rhymes, tone-rhyme combination), probabilities were estimated by marginalizing over the set of character C*_φ_* that share the relevant feature *φ*:

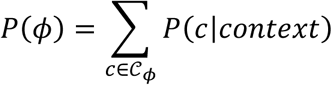

To account for the sequential nature of these units, the summed probabilities were normalized by the probability of the preceding prefix, 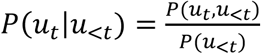, ensuring that the probability reflects the incremental information gain at each linguistic boundary. At all levels, only the top *k* predictions whose cumulative probability reached at least 0.9 were taken into consideration, thereby preserving the main probability mass while excluding the low-probability tail of the distribution. In addition, *k* was set to a minimum of 40 to ensure sufficient candidate information even in low-entropy contexts (Heilbron et al., 2022). Surprisal for each unit was computed as the negative log probability (*I* = − *log*_2_ *P*).

#### Regressors of no interest

To ensure that the observed effects were not confounded by low-level acoustic or other factors, we included several control variables known to drive brain responses: broadband envelope, spectrogram, acoustic edges, pitch, and character frequency. The broadband amplitude envelope was calculated as the magnitude of the analytic signal obtained from the raw speech signal and its Hilbert transform (Di Liberto et al., 2015), and was then downsampled to 120 Hz after applying a zero-phase anti-aliasing filter. The spectrogram was derived by filtering the speech stimuli into 8 frequency bands between 250 Hz and 8 kHz according to Greenwoord’s equation (Greenwood, 1961), then computing the amplitude envelope within each band as before. Including both the envelope and spectrogram enables the model to account for spectral and temporal variations in the acoustic signal (Di Liberto et al., 2015). To capture transient changes in amplitude (acoustic edges; Daube et al., 2019), we included the standard deviation of each phoneme’s amplitude from onset to offset, based on boundaries defined by the MFA. Pitch was calculated for each phoneme using Praat via the parselmouth interface, as prosodic variation has been shown to modulate prediction-related neural responses (Zhang et al., 2021). All acoustic control variables were modeled as impulses aligned with phoneme onsets. Finally, to control for character frequency, we included the unigram surprisal of each character, defined as − log_2_ *P*_(_*_character_*_)_ based on its frequency of occurrence in SUBTLEX-CH (Cai & Brysbaert, 2010).

### Regression analysis

To analyze the mapping between the various linguistic features and the recorded MEG signal, we used multiple temporal response functions (mTRF)—a method widely adopted in naturalistic language research (Crosse et al., 2016). Because speech unfolds rapidly (e.g., characters occur on average every 200 ms), neural responses to adjacent events overlap in time. The mTRF framework addresses this by estimating the contribution of each linguistic regressor across a range of time lags, thereby disentangling overlapping responses. Conceptually, a TRF can be viewed as a linear filter that describes how the brain transforms a stimulus feature, S(t), into a continuous neural response R(t) in the following manner: R(t) = TRF*S(t), where * represents the convolution operator.

All predictors were temporally expanded into time-lagged regressors to model the evoked MEG response. Each lag was assigned a weight, capturing the contribution of that predictor to the brain response at that particular latency. To stabilize estimates in the presence of multicollinearity (both between predictors and across their lags), we applied ridge regression, which penalizes large weights. This allows the model to isolate distinct, time-resolved contribution of each predictor, even when their neural effects overlap in time. In this way, mTRF provides a continuous estimate of neural coding, analogous to an event-related field (ERF/ERP), but with the crucial advantage of explicitly accounting for the temporal overlap and confounds that traditional ERF analyses struggle with in continuous naturalistic input.

The lag range of TRF estimate was initially set from 200 ms before to 1200 ms after event onset (Heilbron et al., 2022), which was then restricted from 100 before event onset to 1000 ms after event onset (-100 ms to 1000 ms) with a step of 1/120 second, as no reliable response was observed outside this range (Di Liberto et al., 2015). All predictors, with the exception of binary onset regressors, were z-transformed prior to model fitting. Correlations among regressors of interest are reported in the Supplementary Materials (Fig. S1).

The MEG data for the mTRF analysis were in the axial gradiometer configuration, which preserved both positive and negative signal polarities. However, because the orientation and location of the underlying neural sources can vary across individuals, averaging the resulting beta coefficients directly across subjects may obscure true effects. To address this, the sensor-level beta coefficients were transformed into planar gradients before averaging. This transformation approximates the local neural response as maximal directly above its neural source, producing positive-only values that are more interpretable and comparable across participants (Hämäläinen et al., 1993). Finally, to visualize the overall temporal profile across sensors, we computed the global field power using only the group-averaged beta coefficients from the temporal sensors. Because the planar-gradient transformation and global field power yield positive-valued response profiles, these profiles do not provide conclusive evidence about the polarity or direction of the underlying deflections. Accordingly, the observed peaks should be interpreted as descriptive characterizations of the response profiles rather than as evidence for sharply separable processing stages.

### Model comparison

Model performance was quantified by calculating the Pearsons correlation coefficients (*r*) between predicted and observed MEG signals. We used a 10-fold cross-validation procedure: for each subject, the 40 trials were randomly partitioned into 10 folds of 4 trials. In each cross-validation iteration, nine folds (36 trials) were used for model training, and the remaining fold (4 trials) served as the held-out test set. Crucially, these train-test partitions were kept identical across all candidate models for each subject; this ensured that any observed differences in predictive accuracy were attributable to the specific predictors rather than variations in trial assignment. For statistical analyses, Pearson’s *r* correlation coefficients were Fisher z-transformed prior to averaging. Averaged values were subsequently transformed back to Peason’s *r* for model comparison and visualization.

### Statistical testing

All statistical tests were two-tailed with an alpha level of 0.05. For within-subject comparisons of model performance, we used paired *t* tests. To compare the amplitude difference between the two onset peaks across all linguistic levels, we calculated for each subject the amplitude difference of the peaks in the first (50-150 ms) and the second (150-300 ms) windows across the four onset types, and employed ANOVA on them. Post-hoc comparisons used Tukey’s honestly significant difference (HSD) test. To compare peak latencies between surprisal, we employed a jackknife-based latency *t* test, which estimates individual peak latencies by iteratively leaving out one participant and recomputing group-level statistics (Miller et al., 1998). To assess whether model performance for surprisal varies linearly with context window size (from 1 to 1024 characters), we fit a linear regression for each participant, with model performance as a function of context window length, and tested whether the resulting slopes differed from zero using a one-sample *t* test.

To examine hemispheric lateralization, we computed laterality index (LI) for each subject, defined as 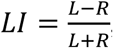, where L and R represent the planar-converted beta coefficients in the left and right temporal sensors, respectively. Positive LI value indicates left-hemisphere lateralization, whereas negative LI value indicates right-hemisphere lateralization. We then tested whether the LI differed from zero using a Bayesian one-sample *t* test (Rouder et al., 2009) implemented in *pingouin* Python package (Vallat, 2018). The Bayesian approach was used because it provides a continuous measure of evidence for or against the null hypothesis (no hemispheric lateralization), enabling more nuanced interpretation and avoiding dichotomous thresholding (Wagenmakers et al., 2018) lateralization. By convention, BF_10_ > 10 indicates strong evidence for lateralization, 3 < BF_10_ < 10 indicates moderate evidence, 1/3 < BF_10_ < 3 indicates anecdotal evidence, and BF_10_ < 1/3 indicates moderate evidence for the absence of lateralization.

## Data availability

All data and code used for stimulus presentation and analysis are freely available at Radboud Data Repository for reviewing purpose and will be made public after acceptance: https://data.ru.nl/login/reviewer-3327085840/NN6FDPQLEF4BYGIXRUOAI63KYE46O5FNK3P4OMY.

## Results

Thirty-four native Mandarin speakers listened to 51 minutes of a Mandarin audiobook while their brain activity was recorded using magnetoencephalography (MEG). Participants were instructed to listen attentively while fixating on a central cross.

To examine whether and how neural responses encoded different linguistic features (e.g., character surprisal), we employed multivariate temporal response functions (mTRF) modeling. We hierarchically added regressors—based on their individual contribution to model fit—and assessed whether each addition yielded a significant improvement, as determined using a cross-validation approach (i.e. the correspondence between model prediction on held-out data and the observed MEG signal; **Fig.1**). In addition to these hierarchical model comparisons, we conducted complementary full-versus-reduced models analyses. Specifically, for each regressor of interest, we compared the full model against a reduced model that included all regressors except the target regressor. This approach allowed us to assess whether a given linguistic feature explained unique variance in neural response beyond that accounted for by the other predictors. We focused our analyses primarily on temporal sensors, given prior evidence implicating temporal regions in language processing (Gagnepain et al., 2012; Willems et al., 2016).

### Onset effect occurs at multiple levels of granularity

We first investigated the neural segmentation of continuous speech across the four levels of granularity: phonemic, sub-syllabic, character, and word levels. In Mandarin, the character serves as the basic unit of the writing system and typically corresponds to a single syllable and a morpheme (the basic unit of meaning in the spoken language). Each syllable is composed of an *initial consonant* (optional), a *final*, and a *tone* (Cheng, 2011; e.g., 她 corresponds to syllable /t-a1/; **Fig. 1**). The final contains one or more vowels, sometimes followed by a nasal consonant. Critically, these sub-syllabic units are the primary perceptual units taught in Chinese reading instruction, whereas phonemes, though the smallest meaning-distinguishing units, are not explicitly emphasized in literacy education. Moreover, while individual characters carry meaning, they are often semantically ambiguous in isolation, and most words are formed by combining two or more characters (Pan et al., 2021; **Fig. 1**).

To control for potential confounds arising from low-level acoustic properties, we first constructed a baseline model incorporating acoustic features, such as the broadband envelope, acoustic edges, pitch, and spectrogram, as impulse regressors (see Methods). We then added regressors of interest: onsets at the phonemic, sub-syllabic, character, and word levels, defined as impulse regressors. Given the inherent temporal overlap and correlation among these onset regressors (Fig. S1), we employed a hierarchical procedure in which the regressor yielding the greatest improvement in model fit was added at each step. This approach allowed us to determine whether each successive level of linguistic structure improved the prediction of neural responses beyond the predictors already included in the model. We further supplemented these analyses with full-versus-reduced model comparisons to test whether each onset regressor explained unique variance beyond all other predictors.

The results showed that the phoneme onset was the most important predictor in capturing neural variance relative to the acoustic baseline model (mean Δ *r* = 0.045, *t*_(33)_ = 17.3, *p* = 4.04 × 10^-18^, *d* = 2.97, 95% CI [0.035, 0.050]; **Fig. 2A**). Adding sub-syllabic onset predictor to this model resulted in a further improvement in fit (mean Δ *r* = 0.003, *t*_(33)_ = 13.3, *p* = 8.33 × 10^-15^, *d* = 2.28, CI [0.0025, 0.0035]; **Fig. 2A**). Word onsets yielded an additional improvement in model performance (mean Δ *r* = 0.0015, *t*_(33)_ = 10.58, *p* = 3.85 × 10^-12^, *d* = 1.81, CI [0.0012, 0.0018]; **Fig. 2A**). In contrast, character onset did not explain significant unique variance beyond the other linguistic levels (mean Δ *r* = 0.0002, *t*_(33)_ = 1.16, *p* = 0.25, *d* = 0.20, CI [-0.0001, 0.0004]; **Fig. 2A**). Complementary full-versus-reduced model analyses yielded a similar pattern, showing that all onset regressors except character onset explained unique variance in the MEG signal (**Fig. S2A-B**). Together, these results suggest that the brain concurrently tracks boundaries across multiple levels of linguistic granularity, from fine-grained phonemic structure to semantically defined word-level representations.

We next investigated the temporal dynamics and spatial profile of these neural responses. Inspecting the global field power of the estimated coefficients from the full TRF onset model across temporal sensors (see Methods), we identified two prominent deflections following each onset type. The first peak, occurring around 100 ms, showed consistent right-hemisphere dominance across all linguistic levels, as quantified by the hemispheric lateralization index (LI, see Methods; phoneme: mean LI = -0.11, BF_10_ = 97.27; sub-syllabic: -0.13, BF_10_ = 612.80; character: -0.11, BF_10_ = 137.48; word: -0.10, BF_10_ = 9.57; **Fig. 2B-C**). The second deflection, peaking around 200 ms, only exhibited right-hemisphere lateralization for character onsets (phoneme: -0.06, BF_10_ = 0.65; sub-syllabic: -0.06, BF_10_ = 0.95; character: -0.09, BF_10_ = 4.42; word: -0.06, BF_10_ = 0.60; **Fig. 2B-C**). Moreover, the relative amplitude of the second deflection, compared to the first, increased systematically from the phoneme to the word level (F_(3, 132)_ = 11.4, *p* = 1.11 × 10^-6^, η^2^ = 0.20; **Fig. 2B**). Post-hoc tests revealed that this amplitude difference was significantly larger for character and word onsets than for phoneme and sub-syllabic onsets (phoneme vs character: *p* = 3.6 × 10^-3^; phoneme vs word: *p* = 1.18 × 10^-5^; sub-syllabic vs character: *p* = 5.7 × 10^-3^; sub-syllabic vs word: *p* = 2.22 × 10^-5^)

**Figure 2.**
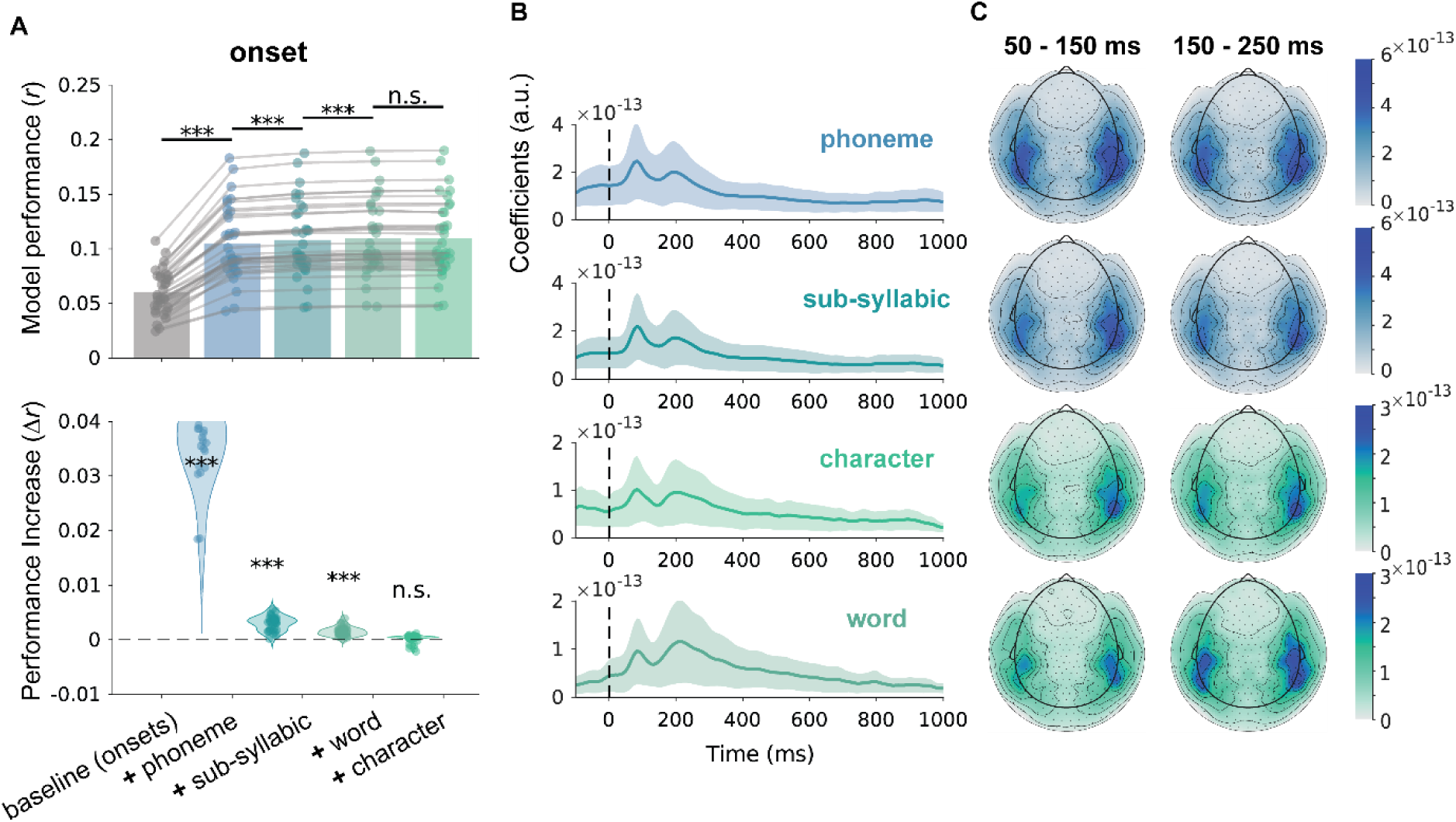
Onset effects at different linguistic levels. **(A)** Model performance when adding regressors with different onset types. The upper panel shows cumulative prediction accuracy for models including different onset regressors. The *baseline* model included acoustic regressors (envelope, spectrogram, acoustic edges, and pitch). Each ‘**+**’ model included one onset regressor on top of the previous model. For example, the ***+*** *sub-syllabic* model contains *acoustics + phoneme onset + sub-syllabic onset*. The order of the added regressors was determined based on their individual contribution to model performance. Dots with connecting lines represent individual participants (averaged over all temporal sensors). The lower panel shows the incremental improvement in model performance (Δ*r*) produced by adding each new onset regressor, providing a clearer visualization of the contribution of each regressor. For instance, the first violin indicates the difference of the +phoneme model compared to the baseline model. **(B)** Temporal profiles of the global field power (GFP) of the beta coefficients from the full onset model (i.e., the ***+*** *character* model), plotted separately for each onset type. Shaded area represents standard deviation. **(C)** Topographic maps of beta coefficients at two peak time windows, shown separately for phoneme, sub-syllabic, character, and word onsets.

### Surprisal effect occurs at multiple levels of granularity

Our finding that Chinese speech is segmented at multiple linguistic levels raises an important question: does language prediction also occur simultaneously across these levels? To address this, we examined prediction at the phoneme, sub-syllabic, character, and word levels.

We quantified linguistic prediction using surprisal, derived from Chinese GPT-2 (Radford et al., 2019; Zhao et al., 2019, 2023), an autoregressive language model trained to predict the next character given prior context. GPT-2 assigns a probability to each predicted token (character) based on its contextual likelihood, from which we computed surprisal (− log_2_(*P*) ) for every character in the audiobook transcript (**Fig. 1**).

To determine the optimal context length, we estimated multiple TRF models using character-level surprisal generated under different context windows (from a single character, up to the model’s maximum of 1024 characters). We found that longer context windows better captured the brain’s response to surprisal (linear fit: *β* = 3.68 × 10⁻⁴, *t*_(33)_ = 12.5, *p* = 2.14 × 10^-14^, **Fig. 3D**), echoing prior behavioral findings that human predictions of upcoming words align more closely with GPT-2’s predictions as context length increases (Goldstein et al., 2022). Accordingly, we used the maximal context window of 1024 characters for all further analyses. Based on these character-level probability estimates, we further derived surprisal measures for other linguistic levels. Word surprisal was calculated from the joint probability of its constituent characters. sub-syllabic surprisal (i.e., initial and final) and phoneme surprisal were calculated as the negative log-transformed probabilities of each unit, obtained by marginalizing across predicted characters sharing the same corresponding phonological features, and, where applicable, conditioned on the probability of the previous prefix (see Methods).

Following the onset-modeling approach, we incrementally added surprisal regressors according to their contribution to model fit, while controlling for acoustic features, all onset regressors, and character frequency, i.e., how frequent the character is in a large text corpus (Cai & Brysbaert, 2010). Frequency was included to ensure that any effects we find are not driven by infrequent characters, but instead truly reflect context-driven surprisal effects (Gillis et al., 2021). In parallel, we conducted full-versus-reduced model comparisons to assess the unique contribution of each surprisal regressor above and beyond all other predictors included in the model.

Our results showed that surprisal significantly improved model performance at all linguistic levels except the phonemic level, corresponding to the finest level of granularity level (**Fig. 3A**). Sub-syllabic surprisal explained the largest amount of additional variance (mean Δ *r* = 4.9 × 10^-3^, *_t_*_(33)_ = 13.5, *p* = 5.46 × 10^-15^, *d* = 2.31, 90% CI [4.1 × 10^-3^, 5.6 × 10^-3^]). Adding character surprisal further improved model fit (mean Δ *r* = 0.8 × 10^-3^, *t*_(33)_ = 11.4, *p* = 1.01 × 10^-12^, *d* = 1.91, CI [0.7 × 10^-3^, 1.0 × 10^-3^]), and word surprisal also explained significant additional variance (mean Δ *r* = 0.1 × 10^-3^, *t*_(33)_ = 2.11, *p* = 0.043, *d* = 0.36, CI [0.005 × 10^-3^, 0.28 × 10^-3^]). In contrast, phoneme surprisal did not improve model performance and instead reduced predictive accuracy (mean Δ *r* = -0.7 × 10^-3^, *t*_(33)_ = -3.78, *p* = 6.27 × 10^-4^, *d* = 0.65, CI [-1.1× 10^-3^, -0.3 × 10^-3^]). Complementary full-versus-reduced model comparisons yielded similar pattern, showing that all surprisal regressors except phoneme surprisal explained unique variance in the neural signal (**Fig. S3A-B**). Together, these findings suggest that predictive processing in Mandarin Chinese operates concurrently across multiple levels of granularity, but not at the finest phonemic level. Because phoneme and sub-syllabic surprisal both reflect phonological prediction and were highly correlated (**Fig S1**), we conducted additional control analyses comparing models containing each regressor separately or both regressors simultaneously. These analyses showed that phoneme surprisal did not explain additional variance beyond sub-syllabic surprisal, whereas sub-syllabic surprisal significantly improved model fit even after controlling for phoneme surprisal (**Fig. S3C**).

The TRF analysis revealed robust N400-like responses, particularly for sub-syllabic (mean latency = 273 ms) and character (mean latency = 309 ms) surprisal. The peak latency for sub-syllabic surprisal was significantly earlier than for character surprisal (jackknife-based latency *t* test: *t*_(33)_ = 3.2, *p* = 2.8 × 10^-3^, *d* = 0.55; **Fig. 3B**). Spatial analyses provided moderate evidence against hemispheric asymmetry across all levels (mean LI = 0.03, 0.02, -0.02, -0.02 for phoneme, sub-syllabic, character and word, respectively; all BF_10_ = 0.2; **Fig. 3C**), indicating bilateral engagement in predictive processing in Mandarin.

**Figure 3.**
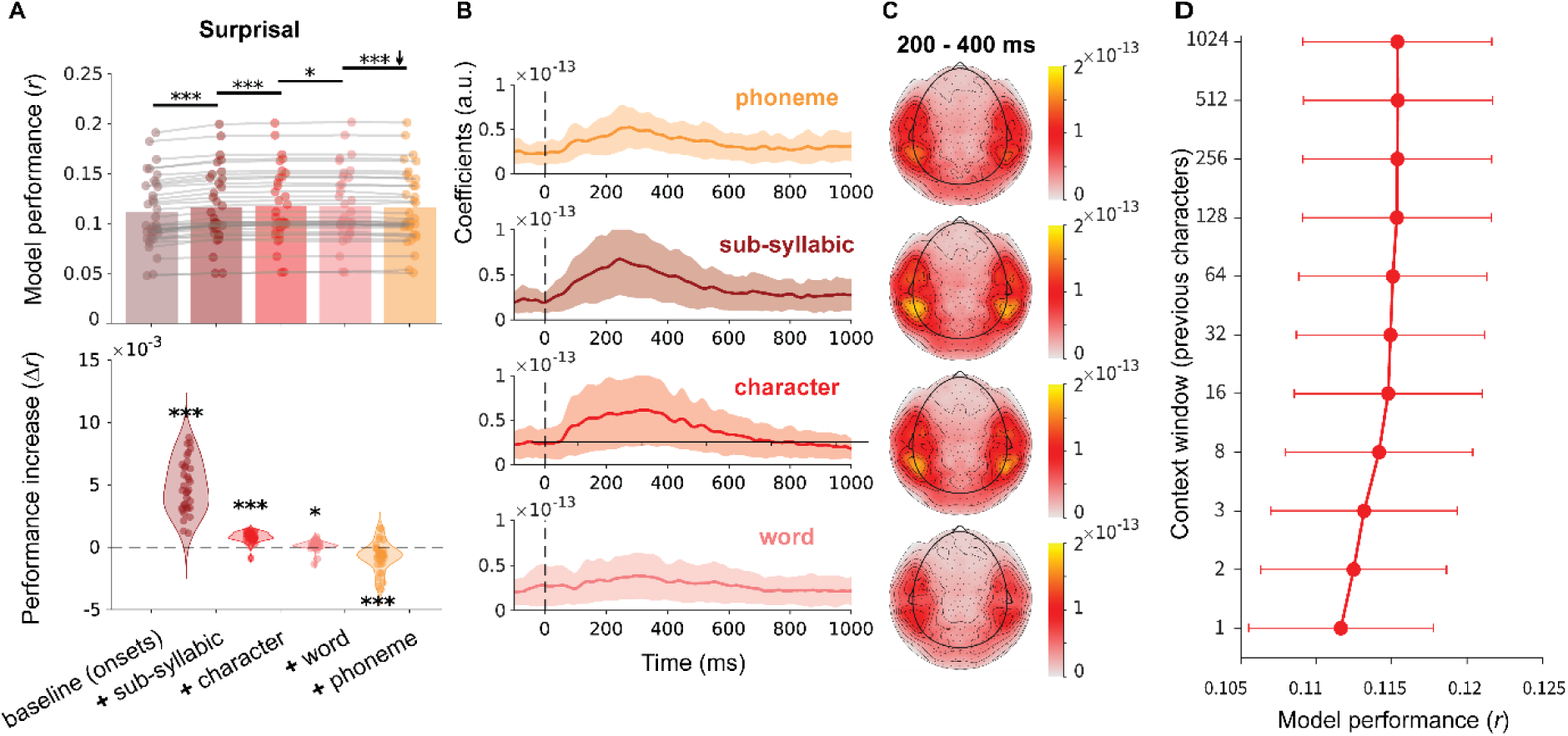
Surprisal effects at different levels of granularity. **(A)** Model performance across different surprisal regressor conditions. The upper panel shows cumulative prediction accuracy for models including different surprisal regressors. The *baseline* (onset) model included acoustic features, all onset regressors, and character frequency. Each ‘***+***’ model included one surprisal regressor on top of the previous model, for instance, the ***+*** *character* model contains *baseline (onset) + sub-syllabic surprisal + character surprisal*. The order of the added regressors is determined based on their individual contribution to model performance. The lower panel shows the incremental improvement in model performance (Δ*r*) produced by adding each new surprisal regressor, providing a clearer visualization of the contribution of each regressor. **(B)** Temporal profiles of the global field power (GFP) of the beta coefficients from the full surprisal model (i.e., ***+*** *word* model), plotted separately for each surprisal regressor. Shaded area represents standard deviation. **(C)** Topographic maps of beta coefficients in the 200-400 ms time window, shown separately for phoneme, sub-syllabic, character, and word surprisal. **(D)** Model performance as a function of context window size (1, 2, 3, 8, 16, 32, 64, 128, 256, 512 or 1024 tokens) used in Chinese GPT-2 to estimate character surprisal, the models included *acoustics + all onsets + character frequency + character surprisal*.

### No independent tonal surprisal effect in Mandarin Chinese

Next, we examined whether listeners generate predictions about lexical tones during speech comprehension. As lexical tones in Mandarin are primarily realized over the rhyme portion of the syllable (i.e. the vowel(s) and any following nasal consonant(s)), we evaluated predictive processing at both the tone and rhyme levels. Tone surprisal and rhyme surprisal were derived from the conditional probabilities of GPT-predicted characters, respectively (see Methods; **Fig. 4A**). Additionally, to assess whether tone and rhyme jointly carry predictive information beyond their individual contributions, we calculated an integrated tone-rhyme surprisal that captures their joint contributions (see Methods; **Fig. 4A**).

We fit TRF models that incorporated tone surprisal, rhyme surprisal, or both, alongside a baseline with the acoustic and onset regressors above. The results showed that adding rhyme surprisal to the model with tone surprisal significantly improved the prediction of neural responses (mean Δ*r* = 1.8 × 10^-3^, *t*_(33)_ = 11.4, *p* = 5.50 × 10^-13^, *d* = 2.31, 90% CI [1.5 × 10^-3^, 2.1 × 10^-3^]; **Figs. 4B**, S3D). Conversely, adding tone surprisal to the rhyme-only model did not yield a significant improvement (mean Δ*r* = -0.01 × 10^-3^*, t*_(33)_ = 0.08, *p* = 0.93, *d* = 0.01, 90% CI [-2.8 × 10^-4^, 2.6 × 10^-4^]; **Figs. 4B, S3D**), suggesting no independent tonal prediction. Crucially, when the integrated tone-rhyme surprisal was added to the model containing both tone and rhyme surprisal additively, model fit improved substantially (mean Δ*r* = 1.0 × 10^-3^, *t*_(33)_ = 13.3, *p* = 7.8 × 10^-15^, *d* = 2.28, 90% CI [8.5 × 10^-4^, 1.2 × 10^-3^]; **Figs. 4B, S3D**). This indicates that the joint contribution of tone and rhyme carries unique predictive information that is not captured by their separate, additive effects. Temporal analyses revealed a neural deflection at approximately 250 ms that was bilaterally distributed (tone: mean LI = 0.02, BF_10_ = 0.3; rhyme: 0.03, BF_10_ = 0.3; tone x rhyme: mean LI = 0.03, BF_10_ = 0.3; **Fig. 4C-D**).

**Figure 4.**
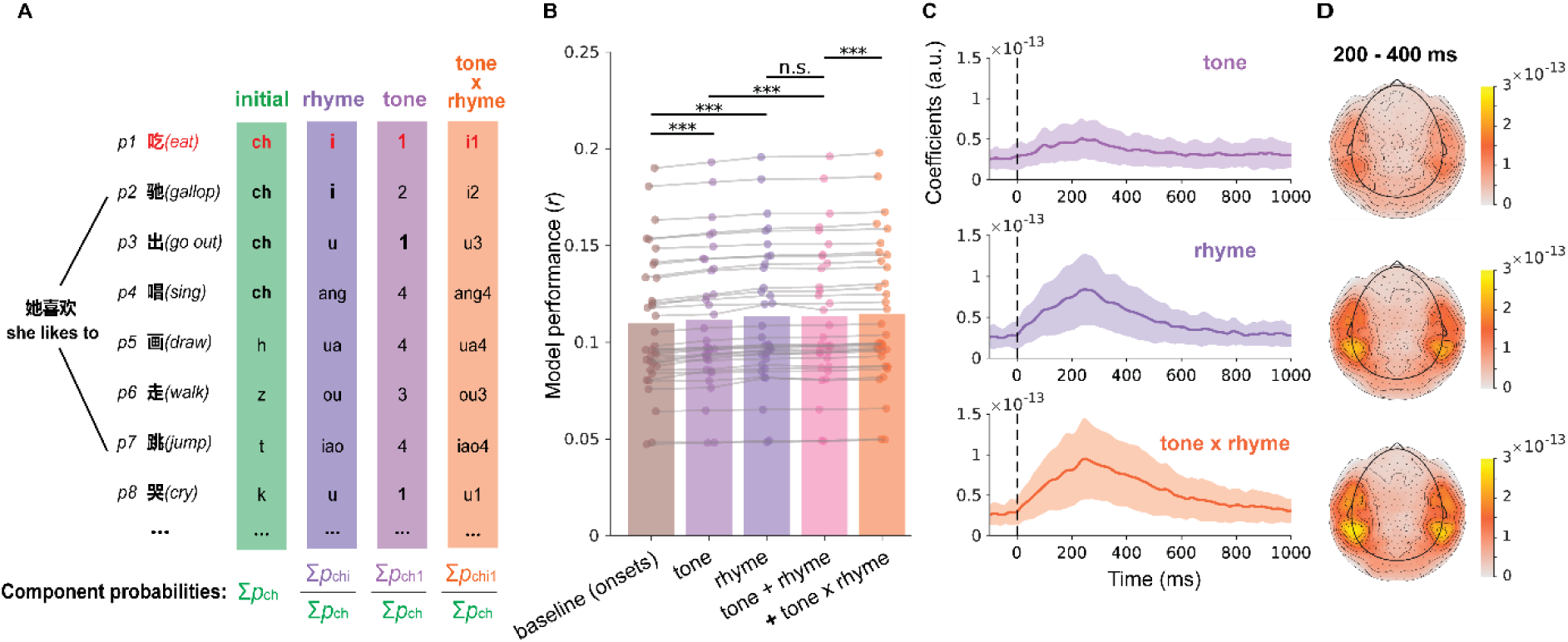
Surprisal effects for tone and rhyme. **(A)** Illustration of probability estimation for different phonological components. GPT-2 generated predictions for upcoming characters with varying probabilities; the target character is marked in red. Each character corresponds to a syllable comprising an initial (green), rhyme (dark purple), and tone (light purple). Initial probability was computed by summing the probabilities of all characters sharing the same initial (e.g., all ‘ch’ characters). Rhyme and tone probabilities were calculated by summing probabilities of characters that share either both the same initial and rhyme (e.g. ch–i) or the same initial and tone (e.g. ch–1), then normalized by the initial probability. Integrated probability was computed by summing probabilities of characters that match the target in initial, rhyme, and tone (e.g. ch–i1), normalized by the probability of the initial. **(B)** Model performance across different surprisal regressor conditions. The *tone + rhyme* model contained tone and rhyme surprisal, whereas ***+*** *tone x rhyme* model included tone surprisal, rhyme surprisal and integrated tone and rhyme surprisal. Shaded area represents standard deviation. **(C)** Temporal profiles of the global field power (GFP) of the beta coefficients from the full model (i.e., the ***+*** *tone x rhyme* model), plotted separately for each surprisal type. **(D)** Topographic maps of beta coefficients in different time windows, shown separately for tone, rhyme and integrated surprisal.

## Discussion

In this study, we examined how different linguistic features modulate brain activity during naturalistic Mandarin speech comprehension by applying linear modelling to MEG signals. Our findings reveal a dissociation between speech segmentation and predictive processing: while the brain tracks linguistic boundaries across a hierarchy of representations, from word down to phoneme levels, prediction (as indexed by surprisal) was most robustly expressed at sub-syllabic, character and word levels, with comparatively limited evidence at the phonemic level. Furthermore, tonal surprisal alone does not strongly modulate neural responses, but contributes when integrated with its phonological base, highlighting the interdependence of segmental (rhyme) and suprasegmental (tone) cues.

### Speech is segmented into different units of granularity

Our first finding is that the listening brain is sensitive to linguistic onsets spanning from phoneme to word, indicating parallel segmentation of continuous speech into units of different sizes. This supports theoretical frameworks proposing that listeners use a hierarchy of cues to segment speech (Brent, 1999; Mattys et al., 2005). Neurophysiological evidence from English has shown that the brain concurrently tracks phoneme and word boundaries (Karunathilake et al., 2023, 2025), even under noisy conditions (Brodbeck et al., 2018). Our data extend these findings to Mandarin by showing parallel tracking from phonemes to words, while also revealing sensitivity to boundaries of language-specific sub-syllabic units, namely initials and finals. This effect may arise from the acoustic and articulatory properties of Mandarin finals: the transitions between phonemes are temporally compressed and coarticulated, such that multiple vowels or nasals are perceived as a cohesive perceptual chunk. This temporal compactness may foster perceptual binding at the sub-syllabic level. Such multi-level segmentation may provide complementary information, facilitating robust and rapid processing in the face of noise and variability (Ding & Simon, 2014; Gross et al., 2013; Keitel et al., 2018; Saberi & Perrott, 1999). In addition, we observed two deflections following all onset types. The first (∼100 ms) resembles the N100, a well-established neural marker of word onset effects in both artificial and naturalistic language (Brodbeck et al., 2018; Sanders et al., 2002; Sanders & Neville, 2003; Tezcan et al., 2023). The second deflection (∼200 ms) exhibited a graded increase in amplitude relative to the first from phoneme and sub-syllabic to character and word levels. This graded pattern likely reflects the integration of semantic information: as characters and words carry morphemic or lexical meaning, binding these features into the evolving discourse model may demand greater computational resources (Broderick et al., 2018, 2019; Huang et al., 2014; Van Den Brink et al., 2001).

### Linguistic prediction occurs in parallel at multiple levels except for phonemic level

Next, our results indicate that neural responses are sensitive to surprisal across multiple levels of granularity, with each level capturing unique context-dependent information. This supports theoretical proposals that predictive processing operates across varying timescales and levels of representational abstraction (Altmann & Mirković, 2009), and extends empirical findings from English to Mandarin (Gillis et al., 2021; Heilbron et al., 2022). Notably, predictive signals were predominantly driven by sub-lexical surprisal, particularly at sub-syllabic and character levels. This pattern aligns with the fundamental incrementality of language comprehension: rather than waiting for an entire lexical unit to unfold, listeners may continuously refine expectations at each sub-lexical step, generating rapid and robust “micro-predictions” (Brodbeck et al., 2018; Donhauser & Baillet, 2020). Such multi-level sensitivity to surprisal may enable efficient resolution of ambiguities and redundancies that arise at different stages of processing, thereby guiding the real-time interpretation of continuous speech with remarkable speed and resilience (Christiansen & Chater, 2016).

A key finding of this study is the absence of a reliable unique surprisal effect at the phoneme level, despite clear neural sensitivity to phoneme boundaries. This dissociation suggests that although the brain segments speech into fine-grained phonemic units, predictive processing rely more strongly on language-specific and functionally salient representational units. In Mandarin, sub-syllabic units constitute the foundational components of literacy instruction and pronunciation training, whereas phonemes are not typically taught as explicit linguistic units. The reduced emphasis on explicit phonemic knowledge may therefore limit the extent to which phoneme-level representations support predictive processing (Huettig et al., 2022). While some neurophysiological studies have indicated that phonemes serve as fundamental units of phonological encoding in Mandarin (Qu et al., 2012, 2020), behavioral evidence suggests phonemes play a subordinate role in word production planning (Chen et al., 2016; O’Seaghdha et al., 2010). Notably, these studies primarily focused on initial phonemes, which overlap substantially with the initial component of sub-syllabic structure, while largely overlooking the role of finals. Evidence from Cantonese, a closely related Chinese dialect, demonstrates that shared finals can drive priming effect (Wong & Chen, 2008, 2009), underscoring the functional relevance of sub-syllabic units. Moreover, sub-syllabic awareness has been shown to be a critical predictor of early reading acquisition in Chinese (Shu et al., 2008; Siok & Fletcher, 2001). From a distributional perspective, higher transitional probabilities within finals relative to those across initial-final boundaries, along with the frequent co-occurrence of phonemes within finals, may further reinforce sub-syllabic units as stable processing units (Hay & Baayen, 2005). Together, these findings suggest that, although phonemes provide a fine-grained articulatory scaffold, sub-syllabic units may constitute a particularly relevant level for prediction-related processing during Mandarin speech comprehension.

### Tone information is not predicted independently in Mandarin

We found that tonal prediction does not operate independently in natural Mandarin speech comprehension, whereas prediction of the rhyme (i.e., the non-initial portion of a syllable) uniquely explains variance of the neural signal. This aligns with previous findings from eye-tracking, which used the visual world paradigm and failed to observe anticipatory fixations toward tonal competitors and reported only marginal effects for rhyme, suggesting that rhyme—but not tone—is pre-activated predictively (Xu et al., 2024; Zhao et al., 2024). In contrast, several ERP studies have shown that explicit tonal violations can elicit N400 effects (Brown-Schmidt & Canseco-Gonzalez, 2004; Hu et al., 2012; Zhao et al., 2011). However, these paradigms often use semantically incongruent or lexically implausible stimuli, which may primarily reflect semantic violation rather than genuine tonal prediction. By quantifying surprisal across all contextually possible continuations weighted by their probabilities, our approach provides a more ecologically valid measure of tonal prediction. Our findings also converge with evidence that rhyme information constitutes a stronger constraint than tone during lexical access (Sereno & Lee, 2015; Wiener & Turnbull, 2016; Yang & Chen, 2022), especially in predictive contexts (Shen et al., 2021).

A recent EEG study reporting decodable tonal predictions (Liu et al., 2025) appears to contradict our null result. We posit that this discrepancy stems from fundamental methodological differences. Their task explicitly required participants to predict one of two highly constrained tonal outcomes, strongly encouraging strategic tonal prediction. In contrast, during naturalistic audiobook listening, the space of possible continuations is vastly larger and less constrained, rendering independent tonal prediction less essential and its neural signature more subtle, and therefore more difficult to detect in aggregate brain responses. Critically, we found that integrated tone-rhyme surprisal provided more predictive power than the sum of its parts. This synergistic effect demonstrates that although tone may not be predicted in isolation, it carries information that is indispensable when combined with its segmental counterpart. This is consistent with behavioral findings showing enhanced anticipatory fixations towards distractors sharing both tone and rhyme with a target (Li et al., 2022; Xu et al., 2024; Zhao et al., 2024). Together, these results suggest that during natural speech comprehension, listeners do not generate independent predictions about tone, but instead integrate tonal and segmental cues to guide efficient interpretation.

### Limitations and future research

While this study provides evidence for prediction-related neural responses at multiple linguistic levels, several limitations should be considered. First, all surprisal measures were derived from character-level GPT-2 next-token probabilities. Although this approach allowed us to estimate predictability across multiple linguistic units in a common probabilistic framework, these measures may not fully capture the computations underlying human predictive processing at each representational level. Ideally, surprisal at each level would be estimated using models specifically trained to predict the corresponding linguistic unit. To our knowledge, however, such models are not currently available for all of the units examined here. Nevertheless, the finding that character-derived sub-syllabic and word surprisal explained unique neural variance beyond character-level surprisal suggests that these measures capture predictive information that is not reducible to character prediction alone. Second, the present analyses were conducted at the sensor level and did not examine the neural sources of the observed effects. Source-level analyses could provide additional insight into the anatomical organization of prediction-related responses across linguistic levels. However, the main goal of the present study was to test whether predictive effects can be observed across multiple levels of linguistic granularity, rather than to localize their precise neural generators. Moreover, although MEG provides useful temporal resolution and some spatial information, closely adjacent neural representations may still be difficult to dissociate reliably, even with source reconstruction. Third, the study was based on a single audiobook. Although the story was naturalistic and the narrator was a native speaker of standard Mandarin, the results may partly depend on material-specific properties, such as vocabulary distribution, prosody, narrator characteristics, and narrative structure. Future studies using more diverse speech materials, multiple speakers, and different discourse contexts will be important for assessing the generalizability of the present findings.

## Conclusions

In summary, our findings support the view that predictive processing is a general feature of language comprehension, while suggesting that its implementation is shaped by language-specific structures. In Mandarin Chinese speech comprehension, prediction does not necessarily engage all phonological sublevels equally; rather, it appears to be most strongly expressed at representational levels that maximize interpretive utility.

## Acknowledgements

This study was supported by China Scholarship Council to QW; European Union’s Horizon Europe research and innovation programme under the Marie Skłodowska-Curie grant agreement for individual fellowship (101106569) and an NWO Veni fellowship to EB; Marie Skłodowska-Curie grant agreement for individual fellowship (101109025) to JF; POLONEZ BIS programme co-funded by the National Science Centre and the European Union’s Horizon 2020 research and innovation programme under the Marie Skłodowska-Curie grant agreement No. 945339 to JS, and European Research Council (ERC Consolidator grant number 101000942) and the Netherlands Organisation for Scientific Research (NWO Vici grant number VI.C.231.043) to FPdL. We thank Jan-Mathijs Schoffelen for advice on the data analysis and thank all the authors of the open source software we used.

## Supplementary figures

**Supplementary Figure 1.**
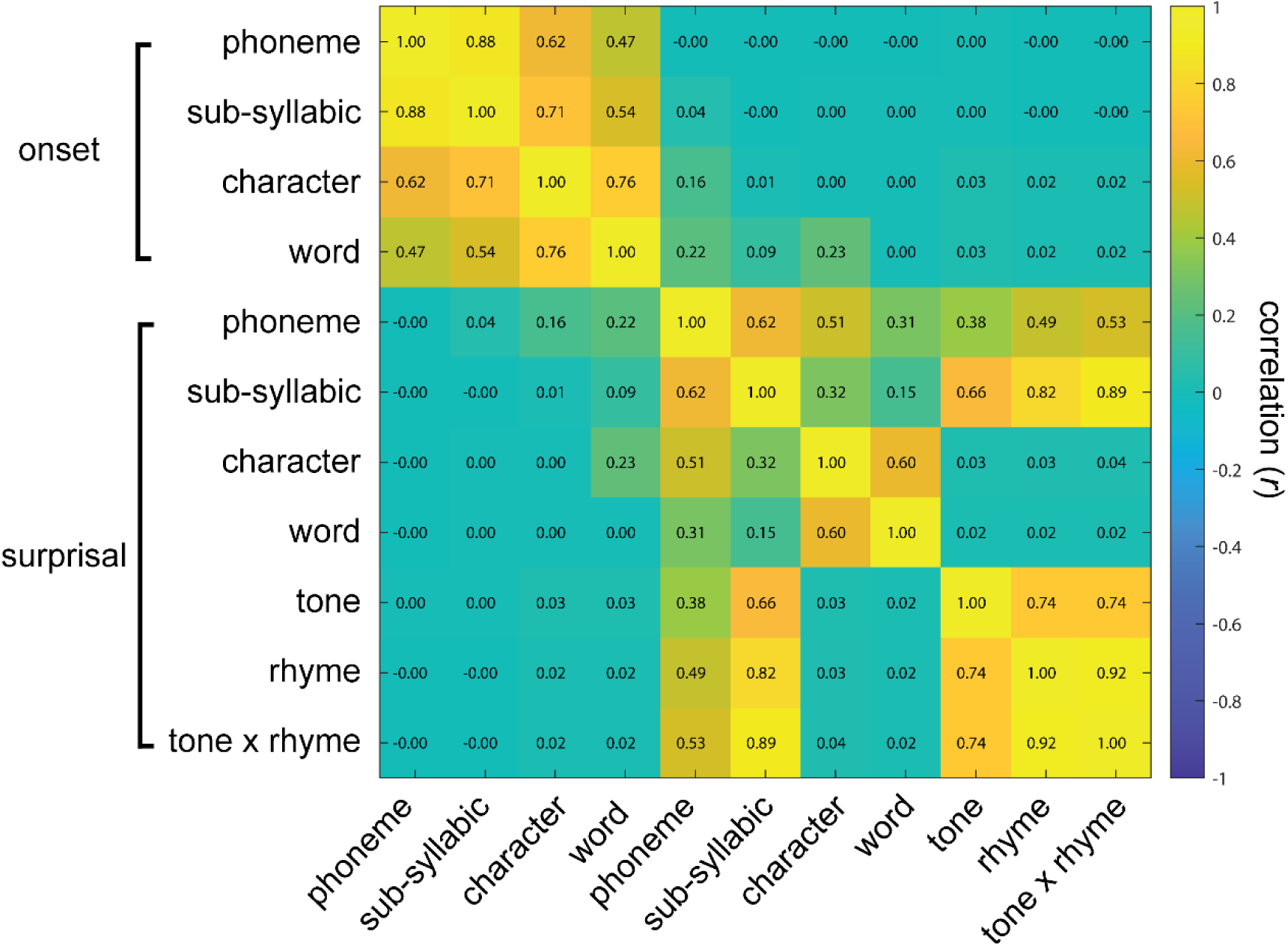
Correlation matrix between regressors. Each cell shows the Pearson’s correlation coefficient between a pair of regressors. Because regressors differ in their linguistic granularity and event rate, correlations were computed after each regressor had been constructed in the format used for temporal response function modelling. Specifically, each regressor was represented as a time-series impulse function spanning the full duration of the neural signal.

**Supplementary Figure 2.**
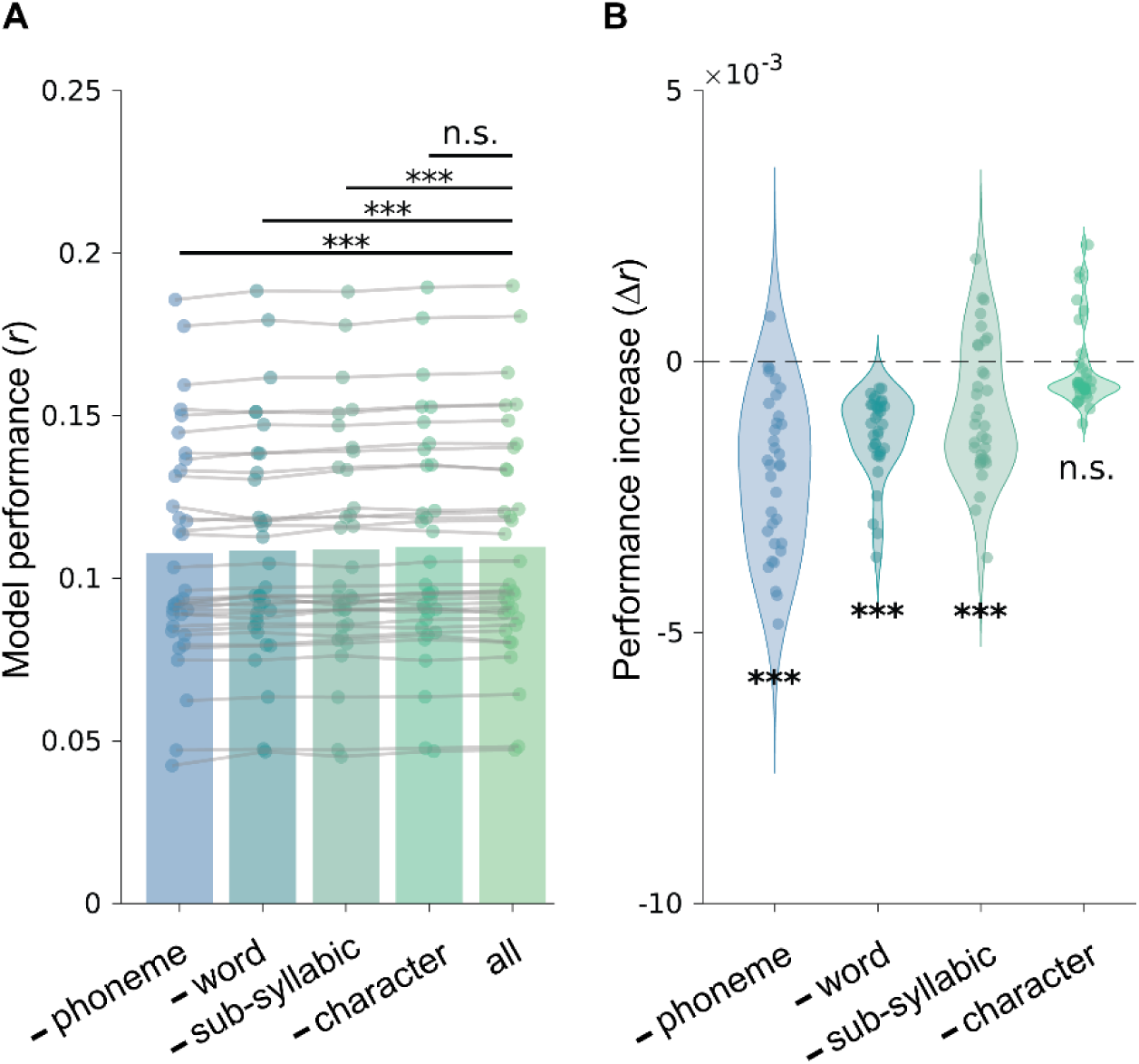
Full-versus-reduced onset model. (A) Model performance across reduced onset models. Bars show the cumulative prediction accuracy for models in which one onset regressors was excluded from the full onset model. Each ‘-’ model excluded one onset regressor from the full model, which contains all onset regressors. Models are ordered according to their average prediction performance. (B) Unique contribution of each onset regressor, quantified as the decrease in model performance (Δ*r*) when that regressor was excluded from the full onset model. A significant decrease in model performance indicates that the excluded regressor explains unique variance in the neural response.

**Supplementary Figure 3.**
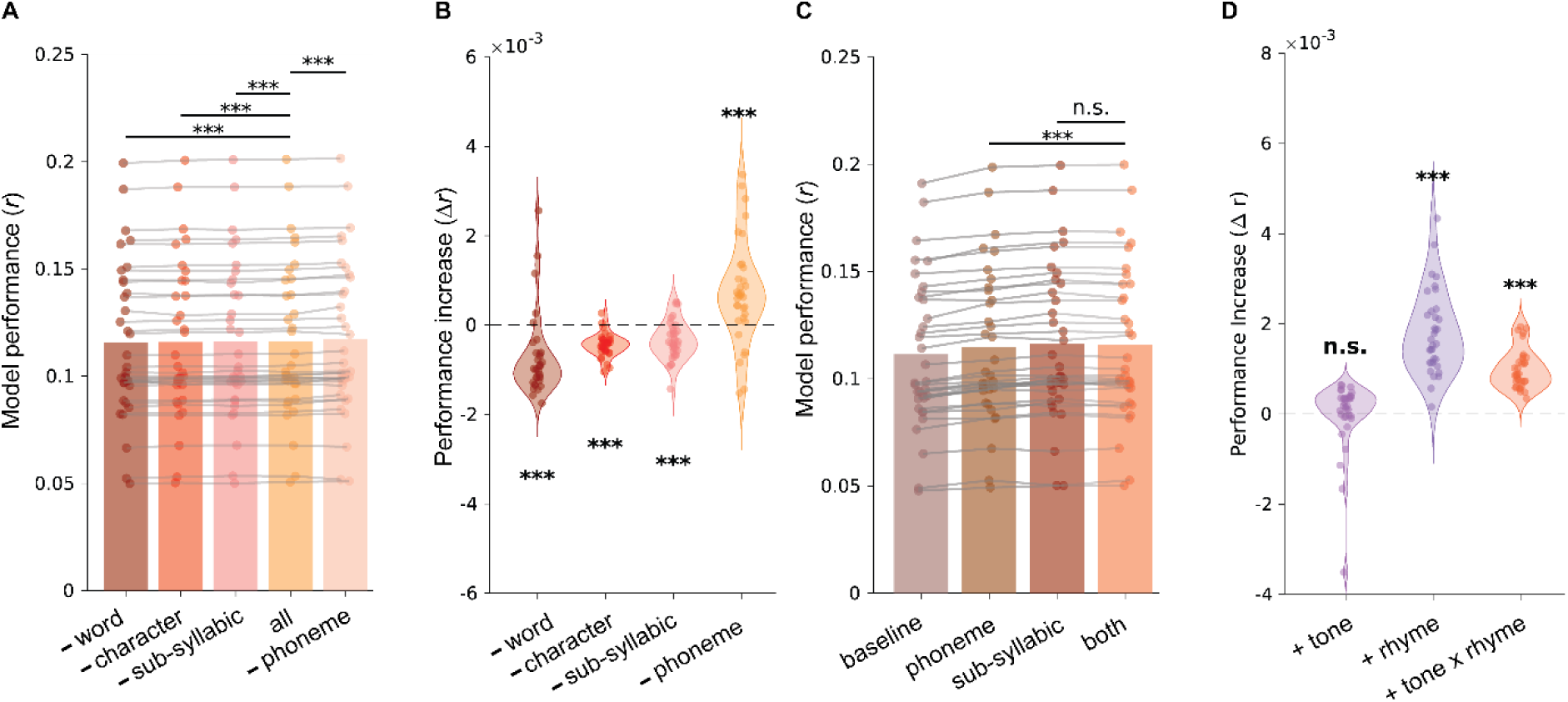
Full-versus-reduced surprisal model comparisons. (A) Model performance across reduced surprisal models. Bars show the cumulative prediction accuracy for models in which one surprisal regressor was excluded from the full surprisal model. Each “−” model excluded one surprisal regressor from the full model, which contained all surprisal regressors. Models are ordered according to their average prediction performance. (B) Unique contribution of each surprisal regressor, quantified as the decrease in model performance (Δ*r*) when that regressor was excluded from the full surprisal model. This provides a clearer visualization of the variance explained by each surprisal regressor beyond the remaining predictors. A significant decrease in model performance indicates that the excluded regressor explains unique variance in the neural response. (C) Comparison between phoneme and sub-syllabic surprisal models. The baseline model contained all acoustic and onset regressors, as well as the lexical frequency regressor. The phoneme model included the baseline model plus phoneme surprisal; the sub-syllabic model included the baseline model plus sub-syllabic surprisal; and the combined model included the baseline model plus both phoneme and sub-syllabic surprisal. Adding sub-syllabic surprisal to the phoneme surprisal model significantly improved model performance (*t_(33)_* = 8.4, *p* = 9.88 × 10^-10^, *d* = 1.44), whereas adding phoneme surprisal to the sub-syllabic surprisal model did not improve model performance and instead showed a marginal decrease (*t_(33)_* = -2.0, *p* = 0.052, *d* = 0.33). These results suggest that phoneme surprisal does not explain unique variance even beyond sub-syllabic surprisal. (D) Incremental improvement in model performance (Δ*r*) for tone, rhyme, and integrated tone-rhyme surprisal. The three violins show, respectively, the contribution of tone surprisal beyond rhyme surprisal, rhyme surprisal beyond tone surprisal, and integrated tone-rhyme surprisal beyond the separate tone and rhyme surprisal regressors.

